# Long-projection astrocytes challenge canonical territorial organization in the sleep-promoting VLPO

**DOI:** 10.64898/2026.03.06.709635

**Authors:** Félix Camille Bellier, Lou Zonca, Quentin Perrenoud, Leana Razaghi, Laura Dumas, Jason Durand, Laure Lecoin, Karine Loulier, David Holcman, Frédéric Chauveau, Nathalie Rouach, Armelle Rancillac

## Abstract

The ventrolateral preoptic nucleus (VLPO) is a key hypothalamic hub for non-rapid eye movement sleep, yet the glial architecture supporting its circuits remains poorly understood. Here, combining genetic labeling, high-resolution imaging and calcium imaging, we uncover unexpected astrocyte diversity in the VLPO. In addition to classical protoplasmic astrocytes, we identify paired “doublet” astrocytes associated with high local proliferative activity, as revealed by EdU incorporation. We further describe a population of long-projection astrocytes extending processes far beyond canonical astrocytic territories and contacting distant cells. These projections challenge the classical territorial organization of astrocytes and resemble morphologies previously thought to be restricted to hominid brains. Notably, VLPO astrocytes display robust spontaneous Ca²⁺ activity and a highly functionally connected network compared to astrocytes in the cortex and hippocampus. Together, these findings reveal specialized astrocyte architectures and enhanced glial network integration within a sleep-promoting nucleus.

**Reporting summary:** Bellier et al. identify three astrocyte subtypes in the sleep-promoting VLPO, including long-projection astrocytes with hominid-like morphology. They uncover marked postnatal gliogenesis, distinctive spontaneous Ca²⁺ dynamics, and tightly interconnected astrocytic networks, revealing region-specific astrocyte specialization and enhanced glial communication.

## Introduction

Astrocytes are active regulators of neuronal circuits, modulating synaptic transmission, extracellular homeostasis and neuronal network activity through different signaling mechanism^1–4^. Through their close proximity to synapses, astrocytes sense neuronal activity via ion channels, neurotransmitter receptors and transporters, and regulate synaptic signaling through mechanisms including neurotransmitter clearance, ionic buffering and the release of neuroactive molecules. Astrocytes can also dynamically adjust the proximity of their processes to synapses, thereby influencing extracellular diffusion and synaptic efficacy^5^. Through these mechanisms, astrocytes regulate synaptic plasticity and neuronal network dynamics.

Astrocytes have also emerged as important regulators of sleep–wake states. Changes in cortical astrocytic intracellular Ca²⁺ signaling precede sleep–wake transitions and contribute to the regulation of non-rapid eye movement (NREM) sleep depth and duration^6–8^. Astrocyte Ca²⁺ activity in other sleep-related brain regions also modulates sleep–wake states^9^. In the ventrolateral preoptic nucleus (VLPO), a hypothalamic structure essential for NREM sleep regulation, optogenetic stimulation of astrocytes increases sleep duration^8^. Astrocytes can also release sleep-regulating molecules that influence sleep homeostasis and brain state regulation^7,10,11^.

Beyond their interactions with neurons, astrocytes are strategically positioned at the interface between neural circuits and the vasculature. Astrocytic endfeet tightly enwrap blood vessels, enabling astrocytes to mediate neurovascular coupling and match local blood flow to neuronal activity and metabolic demand^12–16^. Astrocytes also contribute to blood–brain barrier integrity and participate in glymphatic exchange processes regulating to the glymphatic wash occurring during sleep^17^.

Despite these pleiotropic functions, astrocytes were historically considered a relatively homogeneous cell population. While neuronal heterogeneity within and across brain regions has long been recognized^18^, including the identification of five neuronal subtypes in the VLPO^19^, astrocytes were traditionally viewed as morphologically and functionally uniform. This view has been revised by recent studies revealing substantial astrocyte diversity across brain regions and developmental stages^20,21^.

In rodents, two principal astrocyte morphologies have been recognized: fibrous astrocytes in white matter and protoplasmic astrocytes in gray matter^22^. Protoplasmic astrocytes typically occupy spatially distinct three-dimensional territories that tile the neuropil with minimal overlap^23^ while remaining interconnected through gap junctions, forming extensive astrocytic communication networks^24^. In humans and higher primates, additional astrocyte morphologies have been described, including interlaminar astrocytes and varicose projection astrocytes, which extend long processes across cortical layers^25–28^. Specialized astrocytic forms are also present in other brain regions, such as Müller glia in the retina and Bergmann glia in the cerebellum^29^. Beyond morphological classifications, recent transcriptomic studies have further revealed substantial molecular heterogeneity of astrocytes across and within brain regions^30,31^.

Despite the emerging role of astrocytes in sleep regulation, the astrocytic organization of the VLPO remains largely unexplored. Characterizing astrocyte diversity in this sleep-promoting nucleus may therefore provide new insights into glial contributions to sleep circuits.

Here we uncover unexpected astrocyte diversity in the VLPO and identify three astrocyte subtypes: protoplasmic, doublet and long-projection astrocytes, distinguished by their morphology and functional properties. These findings reveal previously unrecognized astrocyte architectures in a key sleep-regulating nucleus and suggest that glial organization in hypothalamic circuits may differ from canonical astrocyte patterns described in cortical regions.

## Methods

### Animals

GFAP-eGFP, *Aldh1l1*-eGFP, MAGIC Markers, Gal-GFP, GFAP-CreERT2, and Gcamp6f, Aldh1l1-EGFP, Aldh1l1CreERT2 mice were previously characterized and here used from both sexes and between postnatal day 4 (P4) and P80. All these mouse lines are on C57Bl6j genetic background, except for the Gal-GFP that are FVB.

The *Aldh1l1*-eGFP reporter mice correspond to the Tg(Aldh1l1-EGFP/Rpl10a)JD133Htz mouse line^32^ enables the labeling of most astrocytes in the brain. In contrast, the GFAP-GFP mouse strain^33^, enables the labeling of only sparse astrocytes. The Aldh1l1-CreERT2 mice correspond to the Tg(Aldh1l1-Cre/ERT2)02Kan mouse line^34^. Transgenic mice designed to trigger multicolor labeling in astrocytes located in a wide range of brain regions were generated in K. Loulier lab. The dissected brains were subsequently sent to the Collège de France for further analysis. These mice were housed in a 12 h light/12 h dark cycle with free access to food, and animal procedures were carried out in accordance with institutional guidelines. Animal protocols were approved by the Languedoc Roussillon animal experimentation ethics committee (CEEACD/N°36). Transgenic mice broadly expressing both CAG-Cytbow and CAG-Nucbow transgenes^35^ (aka MAGIC Markers mice), modified to achieve random expression of mTurquoise2/mCerulean, mEYFP, tdTomato, or mCherry from the broadly active CAG promoter following Cre recombination, were crossed with Aldh1l1-Cre/ERT2 (*B6;FVB-Tg(Aldh1l1-Cre/ERT2)1Khakh*) animals^34^, and their *Aldh1l1-Cre/ERT2;CAG-Cytbow;CAG-Nucbow* offspring analyzed at P30 following a single subcutaneous injection of Tamoxifen (0.2 mg; Sigma, T5648) at P4.

The Gal-GFP mouse strain used for this research project, STOCK Tg(Gal-EGFP)HX109Gsat/Mmucd, identification number 016342-UCD, was obtained from the MMRRC, an NIH-funded strain repository, and was donated to the MMRRC by the NINDS-funded GENSAT BAC transgenic project^36^.

The GFAP-CreERT2::GCaMP6f mice were used for biphotonic calcium imagery. This mouse line expresses the GCaMP6f calcium indicator in astrocytes and was generated by crossing the GFAP-CreERT2 line with the Cre-dependent B6J.Cg-Gt(ROSA)26Sortm95.1(CAG-GCaMP6f)Hze/MwarJ transgenic mouse line (referred as Ai95(RCL-GCaMP6f)-D (C57BL/6J)), expressing a floxed-STOP cassette blocking transcription of the fast calcium indicator GCaMP6f (The Jackson Laboratory, Stock No: 028865)^37^. The GFAP-CreERT2 mice express Cre recombinase (Cre), fused to a mutant form of the estrogen receptor ERT2, which requires tamoxifen to be active, in GFAP-positive cells only. To induce GCaMP6f expression, GFAP-CreERT2::GCaMP6f mice were injected intraperitoneally with tamoxifen (100 mg/kg/day for 3 days).

Mice were housed in temperature-controlled (20–22°C) and light-tight ventilated cabinets, under a 12h light-dark cycle with *ad libitum* access to food and water. The beginning of the day (lights-on, rest phase) was at 9 a.m. (Zeitgeber time 0; ZT-0). The beginning of the night (lights-off, active phase) was at 9 p.m. (ZT-12).

All animals were sacrificed at ZT-2 following procedures that were conducted in strict compliance with our institutional protocols and approved by the European Community Council Directive of 1 January 2013 (2010/63/EU) and the local ethics committee (C2EA-59, ‘Paris Centre et Sud’) and local guidelines for the ethical treatment of animal care (Center for Interdisciplinary Research in Biology in College de France, France). The number of animals used in our study was kept to a minimum.

### Immunohistochemistry and Confocal Microscopy

Mice were deeply anesthetized with Euthasol and perfused intracardially with paraformaldehyde (PFA 2%). The brains were carefully dissected and postfixed overnight with PFA 2% at 4°C. The next day, PFA was removed, and the brains were cryoprotected overnight in PBS containing 30% sucrose. The brains were cut into 80 µm sections with a freezing microtome (Leica, Germany). Slices were stored in PBS at 4°C until use. For immunohistochemistry, slices were blocked with PBS-Gelatin-Triton (PBS with 0.2% gelatin and 0.25% Triton), for 1 h at room temperature, incubated overnight at 4°C with primary antibodies (Table 1) in the blocking solution. Then washed for 1 h at room temperature with PBS-Gelatin-Triton, and incubated for 2 h with secondary antibodies (Table 2) and DAPI (1:1000) in the blocking solution and washed three times with PBS before mounting with Fluoromount (Invitrogen).

**Table 1:** List of primary and secondary antibodies used.

| Primary antibody | Dilution | Species | Supplier | Reference |
| --- | --- | --- | --- | --- |
| Anti-CD44 | 1:200 | Rat | Abcam | ab119348 |
| Anti-Cx30 | 1/500 | Mouse | Invitrogen | 332500, |
| Anti-GFAP | 1:500 | Chicken, | Abcam | ab50738 |
| Anti-GFP | 1:500 | Chicken, | Abcam | ab13970, |
| Anti-GLAST | 1/500 | Rabbit | Frontier Institute | Af660-1 |
| Anti-Kir4.1 | 1:100 | Rabbit | Alomone Labs | APC-035 |
| Anti-IP3R1 | 1/100 | Mouse | Thermo Fischer | P11881 |
| Anti-IP3R2 | 1/500 | Rabbit | Alomone labs | ACC-116 |
| Anti-Laminin | 1/30 | Rabbit | Sigma | L9393-2ML |
| Secondary antibody | Dilution | Species | Supplier | Reference |
| Anti-rat Alexa Fluor555 | 1:1000 | Rabbit | Life Technologies | A21428, |
| Anti-mouse 647 | 1:1000 | Goat | Life Technologies | A21235 |
| Anti-chicken 488 | 1:2000 | Goat | Invitrogen | A11038 |
| Anti-chicken 488 | 1:2000 | Goat | Invitrogen | A11038 |
| Anti rabbit 555 | 1:1000 | Donkey | Life Technologies | A31572 |
| Anti rabbit 555 | 1:1000 | Goat | ThermoFisher | A21429 |
| Anti-mouse Alexa Fluor 555 | 1:1000 | Goat | Life Technologies | A21424 |
| Anti-rabbit Alexa Fluor 647 | 1:1000 | Goat | Life Technologies | A21244 |
| Anti-Rabbit 555 | 1:1000 | Goat | ThermoFisher | A21429 |

Isolated astrocytes were selected based on their fluorescent staining. Z-stack images were acquired at a confocal microscope with a 63X magnification, 0.2 steps (Spinning-disk W1 Zeiss Axio Observer Z1). VLPO astrocytes were selected to ensure an approximately equal number of reconstructed astrocytes for each type. Astrocytes from the cortical region were chosen from layer II/II, while those from the CA1 region of the hippocampus (HPC) were selected from the oriens radiatum.

### Astrocyte Morphological Analysis

GFAP-GFP mice of P50-60 were perfused, cut, and their GFP labeling was enhanced as described above. Images of individual astrocytes were taken with a confocal microscope, reconstructed using the Imaris software (v.9.7.5, Bitplane, Oxford Instrument), and then analyzed. For each cell, we performed a Sholl analysis to evaluate the complexity of branching structures. This technique involves drawing a series of concentric spheres centered on the soma at 5 µm intervals and counting the number of intersections between these spheres and the cell’s branching processes. This analysis provides critical insights into the cell’s maximal branching complexity and the Schoenen ramification index^38^, a metric defined as the ratio between this maximal number of distal intersections and the number of proximal intersections occurring at the 5 µm radius. This index provides a quantitative measure of the extent and efficiency of cellular arborization.

The Sholl analysis also allowed the determination of the Area Under the Curve (AUC) using GraphPad (v.8, Dotmatics). Additionally, Imaris extracted the convex hull, representing the smallest convex polygon that encompasses all the reconstructed astrocytes, excluding any long projection processes. We extracted the total filament length, which is the cumulative length of all the astrocyte’s processes, and the maximal filament branch level, defined as the maximum number of branches measured from the soma to the extremity of the astrocyte. Finally, we measured the volume of the somata and the distances from the soma to the closest border of the astrocyte domain.

### Ward’s Clustering and Silhouette

Unsupervised clustering was performed on 8 morphological parameters: the convex hull, the AUC of the Sholl analysis, the maximum Sholl Complexity, the total filament length, the volume of the soma, the distance from the soma to the closest border of the astrocyte domain, the observation or not of a doublet structure (two adjacent astrocytes with somata less than 6 µm apart) and the observation or not of one or more long-process. These data were standardized by centering and reducing all of the values of the 50 astrocytes using Statistica 6 software (Statsoft, Tulsa, OK, USA). We built the scree plot of eigenvalues and PCA components and selected the first four components, which explain more than 90% of the variance (Figure 1A). We then conducted a cluster analysis using unsupervised clustering with Ward’s method^39^, which does not require predefining the number of classes to be characterized. The clusters generated by Ward’s method were refined using the K-means algorithm in MATLAB (MathWorks). To quantitatively assess the quality of a clustering, we performed a silhouette analysis of the K-means clustering results. A positive silhouette value indicates that, on average, an astrocyte is closer to astrocytes within of its own cluster than to those in other clusters within the parameter space. Conversely, a negative value suggests a potential misclassification.

**Figure 1.**
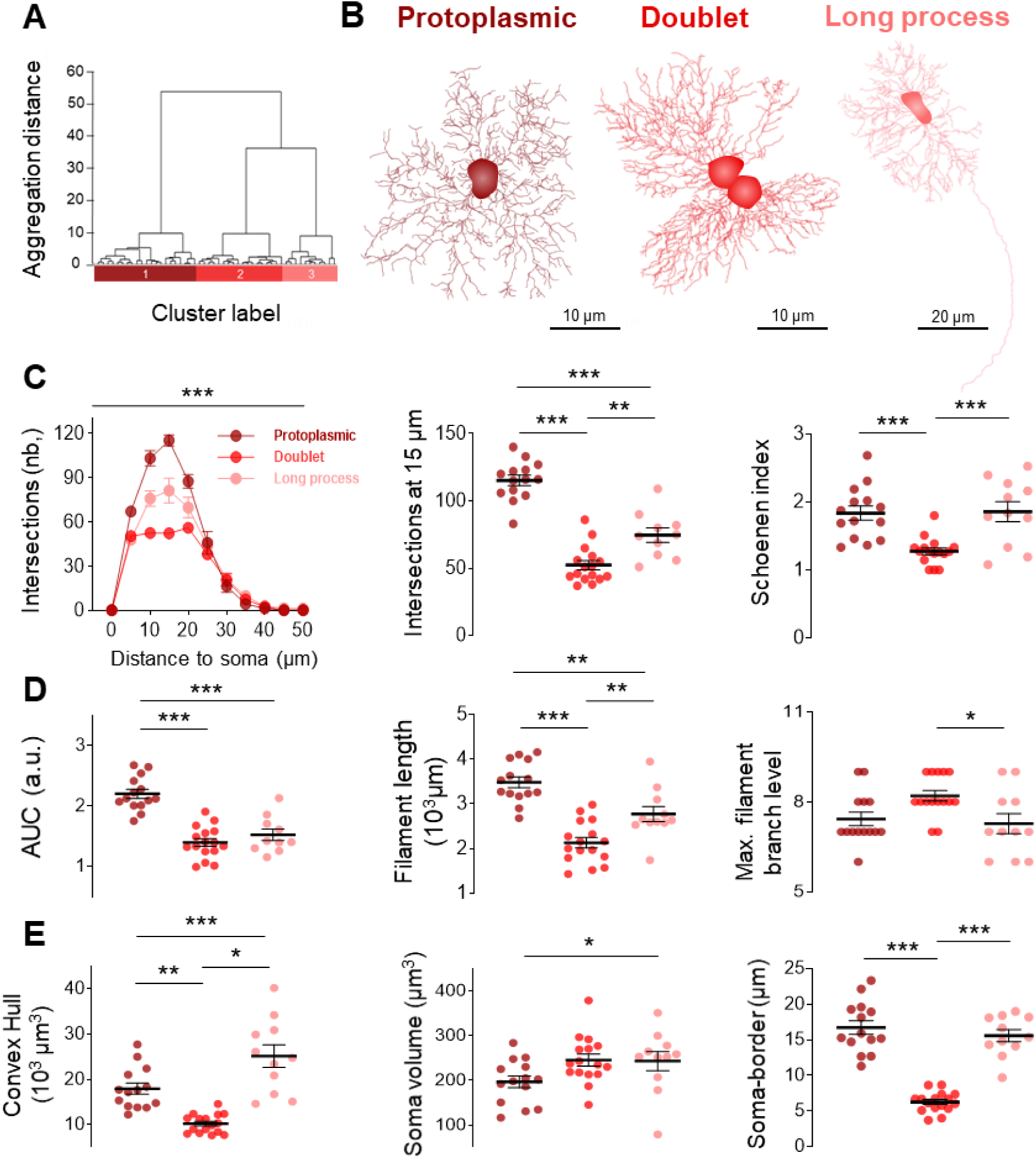
Three astrocyte subtypes in the VLPO with distinct and specific morphological features. (A) Ward’s clustering of 50 astrocytes. Individual cells are represented along the x-axis. The y-axis represents the average within-cluster linkage distance in a space of 4 principal components. Three clusters were identified. (B) Imaris reconstructions of typical astrocytes from Cluster 1 (left), 2 (middle) and 3 (right). (C) Sholl analysis (left), the number of intersections at 15 µm in the Sholl analysis (middle), and their Schoenen complexity index (right). (D) Area under the curve of the Sholl analysis (left), filament length (middle), and maximal branch level (right) of the 3 astrocyte types. (E) Convex Hull (left), soma volume (middle), and the soma-border distances (right). For protoplasmic astrocytes, n=14, from 12 ROIs and 5 animals. For doublet astrocytes, n=16, from 8 ROIs and 5 animals. For long process astrocytes, n=11, from 11 ROIs and 5 animals. Two-way and one-way ANOVA, Kruskal-Wallis, and Tukey’s multiple comparisons test. ***P < 0.001, **P < 0.01, *P < 0.05.

### Corrected Integrated Intensity Analysis

ROIs were manually delineated to quantify the expression levels of various target proteins within astrocytes. We then used the plugin “Proteins Segmentation” (Orion, CIRB), to extract the mean intensity of signal in the ROIs following background subtraction. Corrected Integrated Intensity = Integrated Density − (Area of ROI × Mean Background Fluorescence)

### Quantification of the Different Astrocytic Types in the VLPO

Confocal microscopy images of astrocytes in the VLPO from P80 GFAP-GFP mice, acquired after GFP-labeling enhancement, at a 25X magnification, were used to count astrocyte types manually. A total of 50 z-stacks (mean of 24 images, spaced by 0.5 µm) were used to determine the proportion of each astrocyte type.

### EdU labeling and analyses

To detect DNA synthesis in proliferating cells, we performed 5-ethynyl-2-deoxyuri-dine (EdU) labeling, followed by fluorescent detection^40^. GFAP-eGFP mice were injected i.p. with ∼50 µL at P10, ∼75 µL at P30, and ∼125 µL at P80 of a 10 mg/mL EdU solution prepared in PBS, delivering a dose of 50 mg/kg^41^. Mouse brains were harvested 48 h post-injection. The mice were then perfused, and their brains were sectioned into 100 µm-thick slices using a microtome. Slides were stained with 10 M Alexa568-azide for 30 min and mounted for confocal microscopy. Doublet astrocytes were manually counted in 1 x 1 mm ROIs on stacks of 150 images, each with a 0.2 µm step size.

### Determination of Long-projection Astrocytes Terminations

Four Gal-GFP mice aged 150 days were perfused, and 60 µm slices (prepared as described above) were immunolabeled for GFP, laminin and GFAP, and counterstained with DAPI. Confocal imaging at 60X magnification was used to track 63 astrocytic long-projections within the VLPO. Four main endings were investigated: galaninergic neurons (GFP), cell bodies nuclei (DAPI), blood vessels (laminin), and astrocytes (GFAP).

### Preparation of Acute Slices

GCaMP6f mice aged 50-60 days were decapitated, and their brains were rapidly extracted and submerged in cold (∼0-4°C) slicing artificial cerebrospinal fluid (aCSF) containing (in mM): 119 NaCl, 2.5 KCl, 2.5 CaCl_2_, 26.2 NaHCO_3_, 1 NaH_2_PO_4_, 1.3 MgSO_4_, 11 D-glucose (pH = 7.35). This solution was saturated with a gas mixture of 95% O_2_ and 5% CO_2_. Coronal brain slices, 400 μm thick, were cut using a vibratome (VT1200S; Leica) around Bregma 0 mm for VLPO and cortical recordings, and around Bregma -1.7 mm for hippocampal recordings. Slices were then transferred to a continuously oxygenated (95% O_2_–5% CO_2_) holding chamber containing aCSF for at least 1 hour before recordings.

### Two-Photon Calcium Imaging and Analyses

Individual brain slices were placed in a submerged recording chamber maintained at 32°C and continuously perfused with oxygenated aCSF. The slices were placed under an upright microscope (Axio Examiner Z1 microscope, Carl Zeiss) equipped with a CCD camera (C 2450, Hamamatsu) for high-resolution imaging. GCaMP6f astrocytes were excited at 920 nm, and fluorescence was detected through a 525/40 emission filter and a 580 nm dichroic mirror. Images were acquired at 3 Hz through a 40x water immersion objective (NA 0.95, Olympus) and pre-processed offline with Slidebook imaging software (3i), with a median filter applied to enhance image quality. Two ROI per slice were imaged for 5 min.

ROIs were then manually delineated to selectively investigate calcium dynamics within the soma, proximal processes or distal processes. Within each ROI, activity masks were created to isolate subregions exhibiting physiologically relevant calcium fluctuations. Within each ROI, activity masks were created to isolate subregions exhibiting physiologically relevant calcium fluctuations. The resulting calcium traces were analyzed using Origin (OriginLab Corporation), from which the mean amplitude and frequency of calcium transients were quantified.

### Connectivity Maps

Calcium spontaneous activity in astrocytes can either be pumped back into the endoplasmic reticulum or diffuse to neighboring astrocytes through gap junctions. These calcium transients generate a characteristic bumped shape in fluorescent time-series data. We used a MATLAB toolbox called AstroNet^42^ to perform their analysis (https://github.com/louzonca/AstroNet). Briefly, the signals were summed and normalized for each ROI for 5 min and analyzed over 30 s period to detect the position of active astrocytes. The images were then filtered and binarized to account for the heterogeneity of the luminous intensity. From each detected astrocyte, calcium signals were extracted, and baseline fluctuations were corrected. Events were detected and segmented into distinct occurrences. A connectivity graph was constructed to map the local astrocyte network by following co-activated paths, with each astrocyte constituting a node of the graph. From this analysis, we identified a highly connected subgraph containing hub astrocytes, defined as nodes connected to at least 60% of the network.

### Statistics

All data are expressed as the mean ± standard error of the mean (SEM) and represented as bar graphs or as plots of individual values with the mean value ± SEM. Statistical significance was assessed using appropriate tests based on data distribution and experimental design. Non-parametric analyses were performed using the Mann-Whitney and Kruskal-Wallis tests followed by Dunn’s multiple comparisons test, while parametric t-test and one-way ANOVA, followed by Tukey’s post-hoc test, were applied when data followed a normal distribution. For Sholl analyses, a 2-way ANOVA followed by Tukey’s multiple comparisons test was used. All statistical tests were conducted using GraphPad Prism, and the specific test applied is indicated in the corresponding figure legends. Significance was set at P < 0.05 and expressed as follows: *P < 0.05, **P < 0.01, ***P < 0.001.

Outliers were detected with the median-absolute-deviation (MAD) criterion: the sample median m was first estimated, the absolute residuals ∣xi−m∣ were then computed for each observation, and their median yielded the MAD, a robust scale estimator^43^. Any observation lying outside the interval m ± 3 MAD was classified as an outlier and excluded from subsequent analyses.

## Results

Cellular heterogeneity is a hallmark of brain organization, reflecting the specialized demands of neural circuits across distinct regions. However, the astrocyte landscape within the VLPO remains largely uncharacterized. To address this gap, we employed a comprehensive multimodal approach, combining high-resolution imaging, molecular profiling, and unsupervised classification, to examine the astrocytic population in the VLPO.

### Unsupervised Structural Classification of VLPO Astrocytes Identifies Three Distinct Subtypes

To access the morphological diversity of astrocytes in the VLPO, we selected 50 GFP-positive astrocytes from GFAP-eGFP mice, chosen to capture the broadest possible spectrum of morphological variation. Using high-resolution Imaris 3D reconstructions, we performed an in-depth volumetric analysis of each cell. This included Sholl analysis, to assess arborization complexity, convex hull, and somatic volume measurements. To assess the relative position of the cell body within its occupied territory, we also measured the minimum distance from the soma to its border. Eight morphological features were extracted from each cell (see Materials and Methods). Four principal components were selected from a principal component analysis performed on the discriminative morphological parameters (Figure S1A). Based on these parameters, we employed unsupervised clustering using Ward’s method^39^ to avoid personal bias and group cells with similar characteristics into distinct subpopulations. A clustering tree was constructed, representing individual astrocytes from its leaves and grouping them into the branched ramifications up to a common root (Figure S1B).

**Figure S1.**
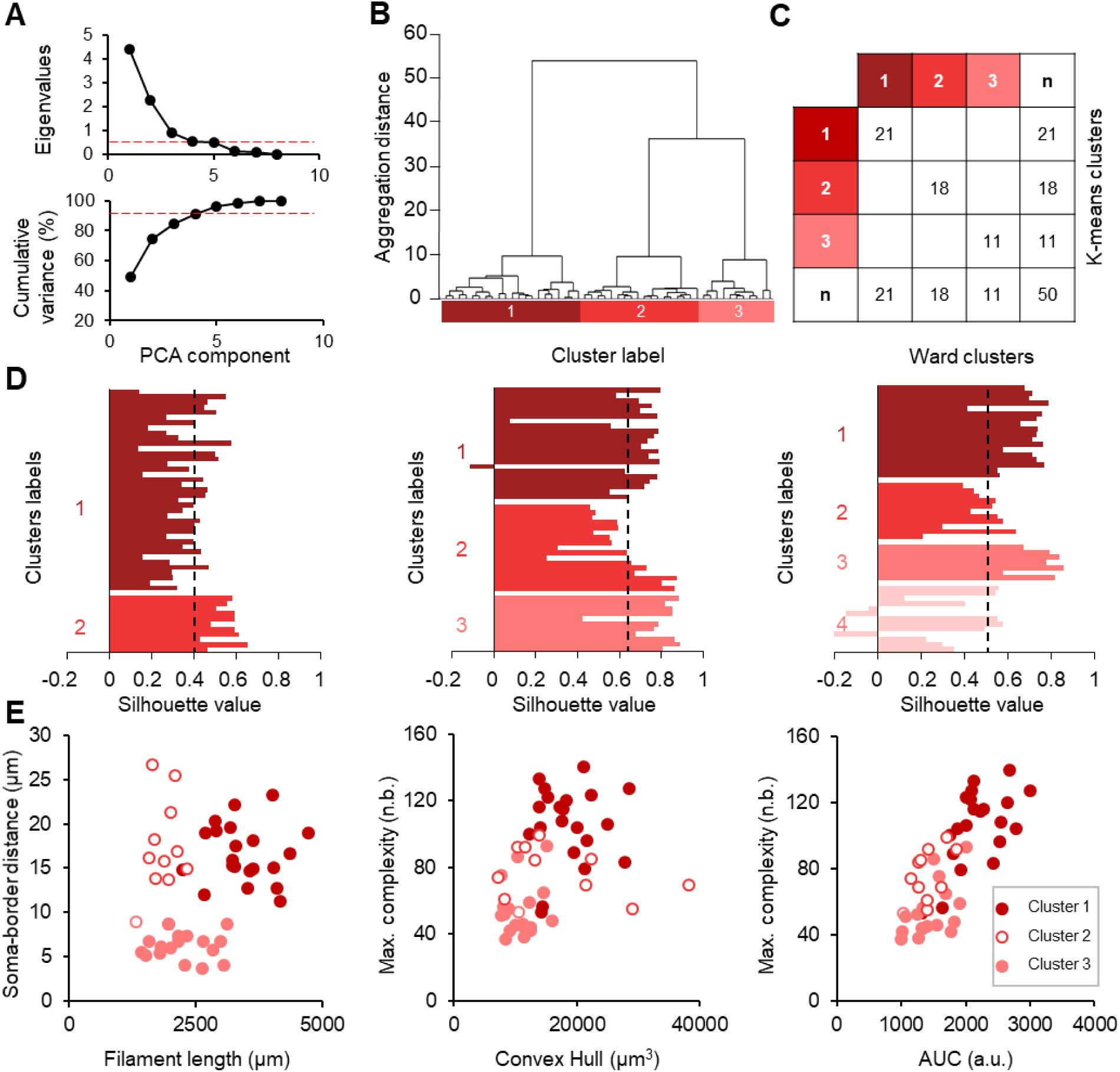
Clustering of VLPO astrocytes based on their morphological properties. (A) Dot plots showing the eigenvalues and the cumulative variance explained by each PCA component extracted from 8 morphological variables. We selected a PCA with 4 components. (B) Ward’s clustering of 50 astrocytes. Individual cells are represented along the x-axis. The y-axis represents the average within-cluster linkage distance in a space of 4 principal components. Three clusters were identified. (C) Clusters generated by Ward’s method (B) were validated by observing the same distribution of astrocytes in the clusters obtained using the K-means algorithm. (D) Silhouette analyses were used to assess the quality of the clustering. The mean silhouette value was 0.4 (dashed line) for 2 Clusters, optimal for 3 Clusters with a mean value of 0.65 (dashed line), and lower for 4 Clusters with a mean value of 0.54 (dashed line). (E) Individual values for each cell among clusters for soma-border distance and filament length (left), among clusters for complexity and convex Hull (middle), and for complexity and AUC (right).

The reliability of Ward’s clustering was validated using the K-mean algorithm. No adjustments were necessary as the K-mean algorithm grouped the cells exactly in the same manner as Ward’s clustering (Figure S1C). To determine whether the allocation of astrocytes into three groups was optimal, we compared our results with clustering trials using fewer or higher groups. We found that a cluster analysis resulting in three groups was the most effective, with a mean silhouette value of 0.65 (Figure S1D). Astrocytes within these three clusters exhibit distinct morphological features (Figure S1E). Consequently, the classification of VLPO astrocytes into three principal subtypes was retained for further analysis.

We next compared the characteristics of these three groups and found that each exhibited highly specific features (Figure 1).

Cluster 1 corresponds to protoplasmic astrocytes. They are the predominant subtype within the VLPO, accounting for ∼71% of the astrocytic population. Astrocytes were initially selected based on their specific characteristics to ensure a sufficient representation of each group for the Ward’s method analysis. A more precise quantification was subsequently performed to assess their proportions within the VLPO. Protoplasmic astrocytes are characterized by a uniform and large distributed arborization around their soma, reflecting their high degree of structural complexity. Sholl analysis reveals that this complexity peaks at 115.21 ± 3.87 intersections at a radial distance of 15 µm from the soma, which is nearly the maximum of all astrocytes (Figure 1B). This complexity is further revealed by their elevated Schoenen complexity index, which normalizes structural intricacy relative to the number of primary processes, providing a refined metric of branching complexity. Consistently, protoplasmic astrocytes exhibit the highest Area Under the Curve (AUC) in Sholl profiles (Figure 1C). Consequently, the cumulative length of their processes is significantly longer than that of other astrocytic subtypes (Figure 1D). Despite their complex arbor and longer filament length level, no increase in filament branching levels was detected (Figure 1D). These astrocytes are also characterized by a small soma, centrally located within their domain (Figure 1E).

Cluster 2, referred to as “doublets”, represents ∼19% of the VLPO astrocytes. These cells are distinguished by the presence of a neighboring astrocyte in extremely close proximity. They have spatially constrained domains and tightly apposed somata (Figure 1A). Consistent with this restricted spatial arrangement, Sholl analysis reveals a markedly reduced morphological complexity, approximately half that in protoplasmic astrocytes, along with a diminished number of intersections at 15 μm and a low Schoenen complexity index (Figure 1B). Doublet astrocytes are also characterized by a reduced area under the Sholl curve and shorter cumulative filament length (Figure 1C), suggesting a compression of their structural development. However, doublet astrocytes exhibit a maximal branch level similar to protoplasmic astrocytes, and even higher than that of long-projection astrocytes, indicating a potential compensatory branching strategy. These cells, therefore appear to have the smaller convex hull volume and the shortest distance from their soma and the nearest boundary of their domain, reinforcing that these cells may have undergone domain compression due to a lack of space (Figure 1D).

Cluster 3 represents ∼10% of VLPO astrocytes and is distinguished by their striking ability to extend one or few processes far beyond the astrocytic domain. These processes are typically straight, unbranched, and can measure >1 mm. Referred to as long-projection astrocytes, these cells are predominantly located near the pial surface and exhibit morphological features in between those of protoplasmic and doublet astrocytes. Accordingly, their overall structural complexity, as assessed by Sholl analysis, falls between the high complexity of protoplasmic astrocytes and the reduced complexity of doublet astrocytes. Despite this intermediate branching profile, they display a similarly low area under the Sholl curve (AUC) to that of doublet astrocytes. This is likely attributable to their sparse distal arborization, leading to a total arborization length that is less extensive than that of protoplasmic astrocytes and more in line with doublet astrocytes. Finally, these astrocytes typically possess relatively large somata, centrally located within their domain (Figure 1D).

### Structural Specialization of Protoplasmic Astrocytes in the VLPO

Protoplasmic astrocytes represent the most abundant astrocyte subtype in the brain^44^. They are located in various brain regions of the gray matter. To investigate whether VLPO protoplasmic astrocytes exhibit region-specific morphological features, we reconstructed astrocytes from two other well-studied brain regions: the cortex and the hippocampus (Figure 2).

**Figure 2.**
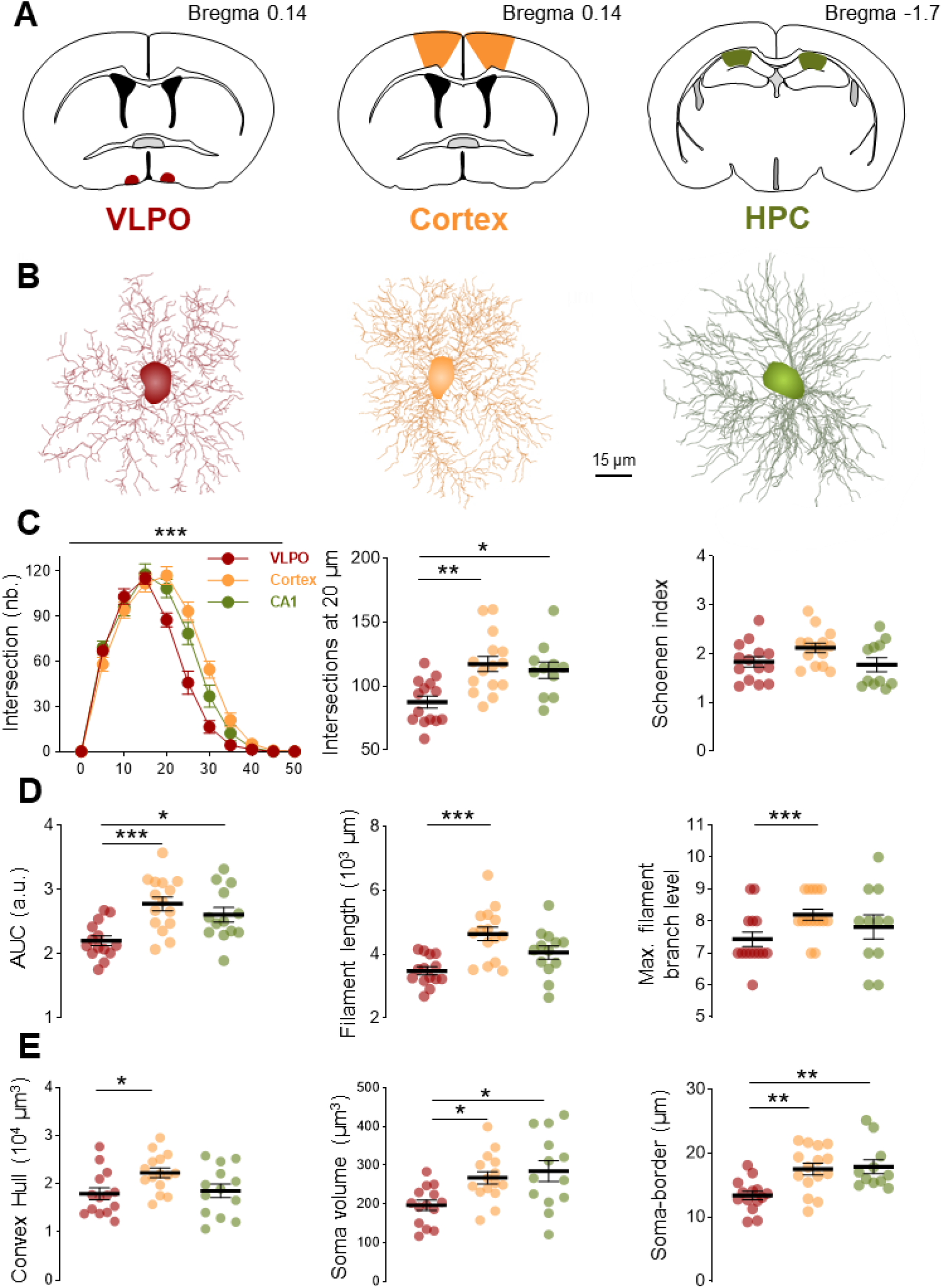
Comparative morphological characterization of protoplasmic astrocytes in the VLPO, cortex, and hippocampus. (A) Anatomical schematics of coronal brain sections showing the localization of the studied brain regions. (B) Morphological reconstructions (Imaris) of typical astrocytes from each region. (C) Sholl analysis (left) of these astrocytes, their complexity at 20 µm (middle), and their Schoenen complexity index (right). (D) The AUC of their Sholl curves (left), their total filament length (middle), and their maximal branch level (right). (E) Morphological analysis of their convex Hull (left), volumes of their soma (middle), and the soma-border distances (right). In the VLPO, n=14, from 12 ROIs and 5 animals. In the cortex, n=15, from 8 ROIs and 5 animals. In the HPC, n=11, from 7 ROIs and 4 animals. Two-way and one-way ANOVA, Kruskal-Wallis, and Dunn’s followed by Tukey’s multiple comparisons test. ***P < 0.001, **P < 0.01, *P < 0.05.

Although VLPO protoplasmic astrocytes exhibit comparable levels of maximal complexity and Schoenen complexity index to those in other regions, their overall arborization is markedly less extensive. This is shown by a significantly reduced arborization complexity, shorter total filament length, lower maximal branch level, and a smaller volume, particularly in comparison to cortical astrocytes. Regarding somatic properties, VLPO astrocytes also display a significantly smaller soma volume relative to both cortical and hippocampus (HPC) astrocytes, and their somata are positioned closer to the borders of their respective domains, consistent with the overall smaller domains of these cells.

### Doublets Correspond to Lately Generated Astrocytes

Although neurogenesis is complete in most regions of the brain at birth, the number of glial cells increases six- to eight-fold during the first three weeks of postnatal development^45^. To assess whether doublet astrocytes, which appear to be subject to spatial constraints in the VLPO, could result from delayed gliogenesis, we first investigated whether they share a common cellular origin. To do so, we used multiplex clonal analysis based on the multicolor MAGIC Markers strategy^46^ to first evaluate whether doublet astrocytes potentially arise from the same pioneer astrocytes. This mouse line enables the visualization of cell lineage, as in this strain, astrocytes derived from the same lineage express identical colors. We thus investigated whether doublet astrocytes consistently display the same color. We found that all doublet astrocytes were sparsely distributed throughout the VLPO and that all the two astrocytes belonging to doublets exhibit the same color, suggesting that they originate from a common progenitor (Figure 4A).

**Figure 3.**
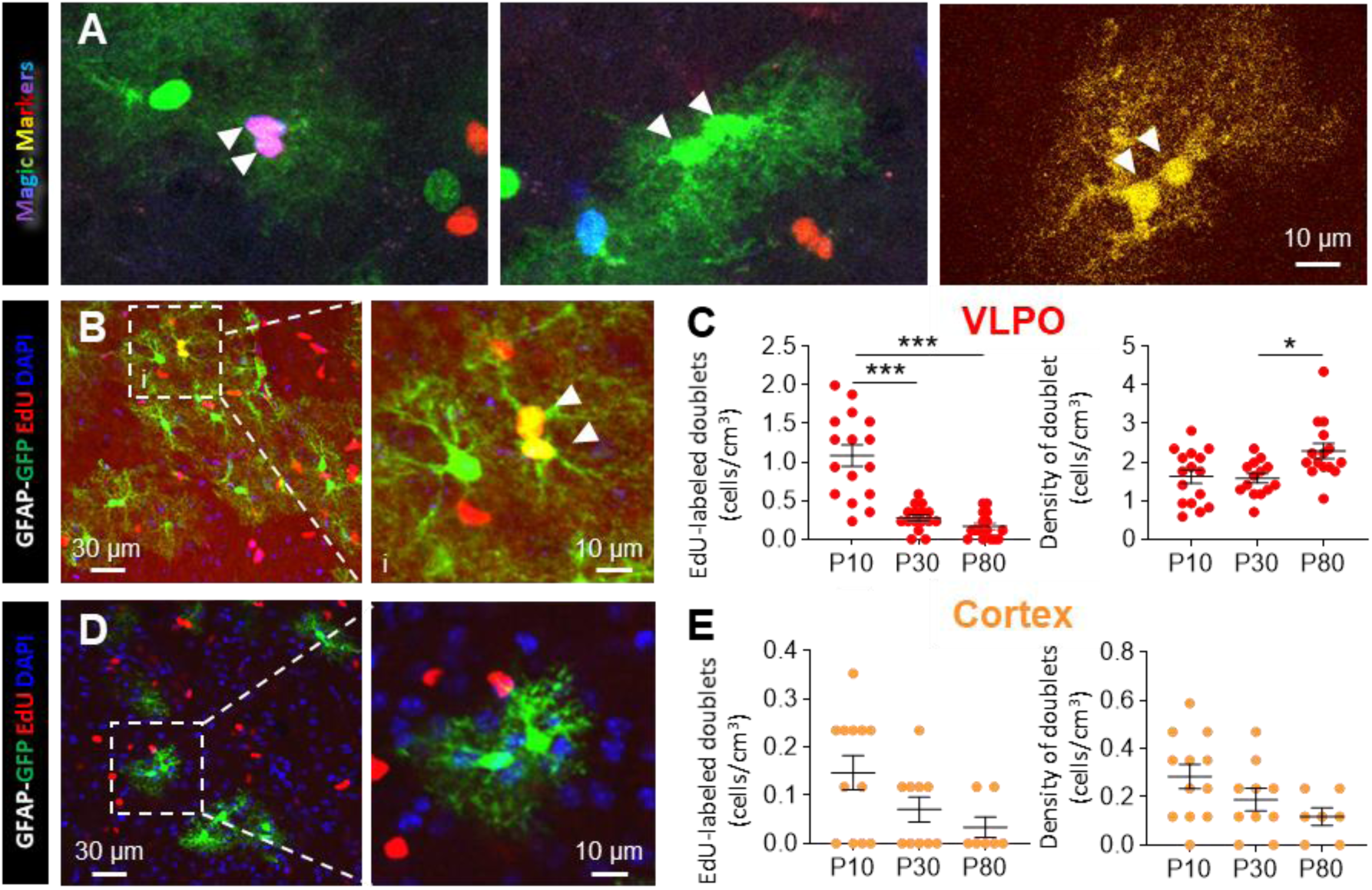
Doublet astrocytes derive from a common progenitor and late gliogenesis in the VLPO. (A) Confocal images of the VLPO from MAGIC Markers mice showing that doublet astrocytes consistently express the same color, indicating a shared origin. Observation on 11 ROIs from 4 mice. (B) Confocal images of EdU labeling in the VLPO (left). Zoom in on a labeled doublet astrocyte (right). Arrowheads indicate the soma of doublet astrocytes. (C) Density of EdU^+^ doublet astrocytes per ROI from 16 ROIs at P10; 18 ROIs at P30; and 17 ROIs at P80 in 4 mice in the VLPO. (D) and (E) Same as in B-C for the cortex from 12 ROIs at P10; 10 ROIs at P30, and 7 ROIs at P80 in 4 mice. Kruskal-Wallis, and Dunn”s multiple comparisons test. ***P < 0.001, **P < 0.01, *P < 0.05.

**Figure 4.**
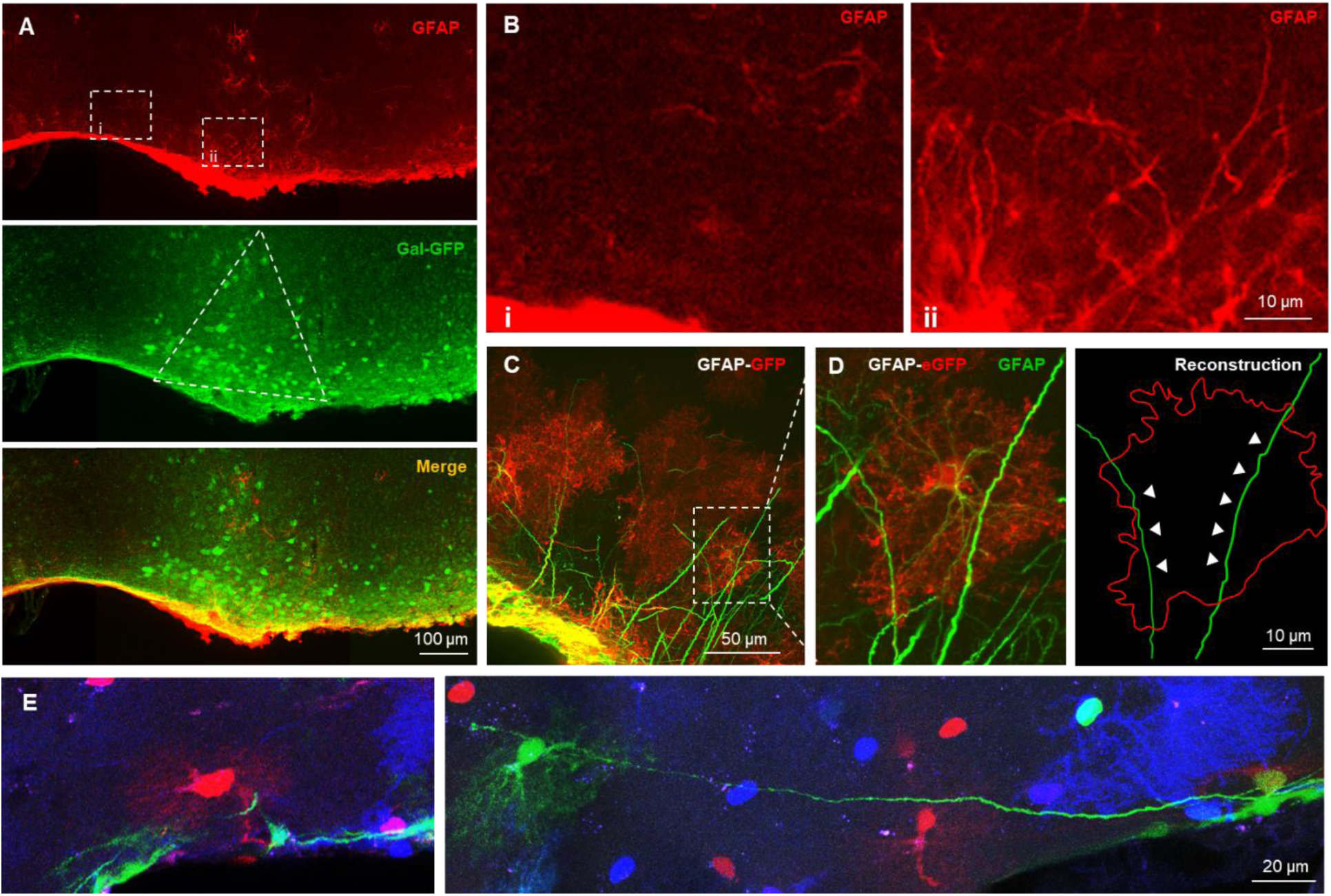
Long-projection astrocytic processes are restricted to the VLPO and cross domains of neighboring astrocytes. (A) Confocal microscopy images of GFAP immunolabeling (red) of the VLPO region of a GFAP-eGFP mouse (green) and the superposition (yellow). (B) Zooms of the ROI indicated by dashed white squares in (A). (C) Confocal microscopy images of GFAP immunolabeling (green) in the VLPO region from a GFAP-eGFP mouse (red) showing a single optical section (z-plane) through the astrocytic domain. (D) Zoom of the white dashed square in (C), highlighting an astrocytic domain intersected by at least two GFAP^+^ processes and reconstruction of these GFAP^+^ processes (green), indicated by white arrowheads, crossing an astrocytic domain outlined by a red line. (E) Confocal images from MAGIC Marker mice showing long projection processes crossing into other astrocytic domains, each marked with distinct colors.

To further explore the greater abundance of doublet astrocytes in the VLPO, we investigated whether they could result from a prolonged mitotic activity during postnatal development. Therefore, we compared astrocyte proliferation dynamics in the VLPO and the cerebral cortex by quantifying newly divided cells through EdU incorporation at three developmental stages: P10, P30, and P80 (Figure 3B–E). This approach enabled temporal tracking of gliogenesis across critical postnatal windows.

At P10, EdU labeling revealed a gliogenesis rate in the VLPO 7 times higher than in the cortex. Although proliferative activity decreased over time in both regions, it remained significantly elevated in the VLPO, about fivefold higher, at P30 and P80. This sustained proliferative rate coincided with a progressive increase in the proportion of astrocytic doublets in the VLPO during postnatal development, in contrast to the cortex, where their frequency steadily declined. By P80, the density of astrocytic doublets in the VLPO was ∼20 times higher than in the cortex (Figure 3B–D), supporting the hypothesis that region-specific postnatal proliferation dynamics contribute to the maintenance and expansion of doublet astrocytes.

To determine whether astrocytic doublets correspond to immature astrocytes, we examined their expression of molecular markers indicative of astrocyte maturation: the glutamate aspartate transporter (GLAST), the inwardly rectifying K(+) channel Kir4.1, and connexin 30 (Cx30), a key astroglial gap junction forming protein (Figure S2). Each of these markers follows a well-characterized developmental expression timeline. GLAST (EAAT1) expression stabilizes between P7 and P21 before declining. Kir4.1 reaches a maximal and stable expression by P21. Cx30 expression initiates at P10 and increases until P50^47^, making it a late-stage maturation marker. At P30, we detected no significant differences in the expression levels of these proteins between astrocytic doublets and protoplasmic astrocytes. These findings suggest that doublet astrocytes are not molecularly immature at this stage (Figure S2).

**Figure S2.**
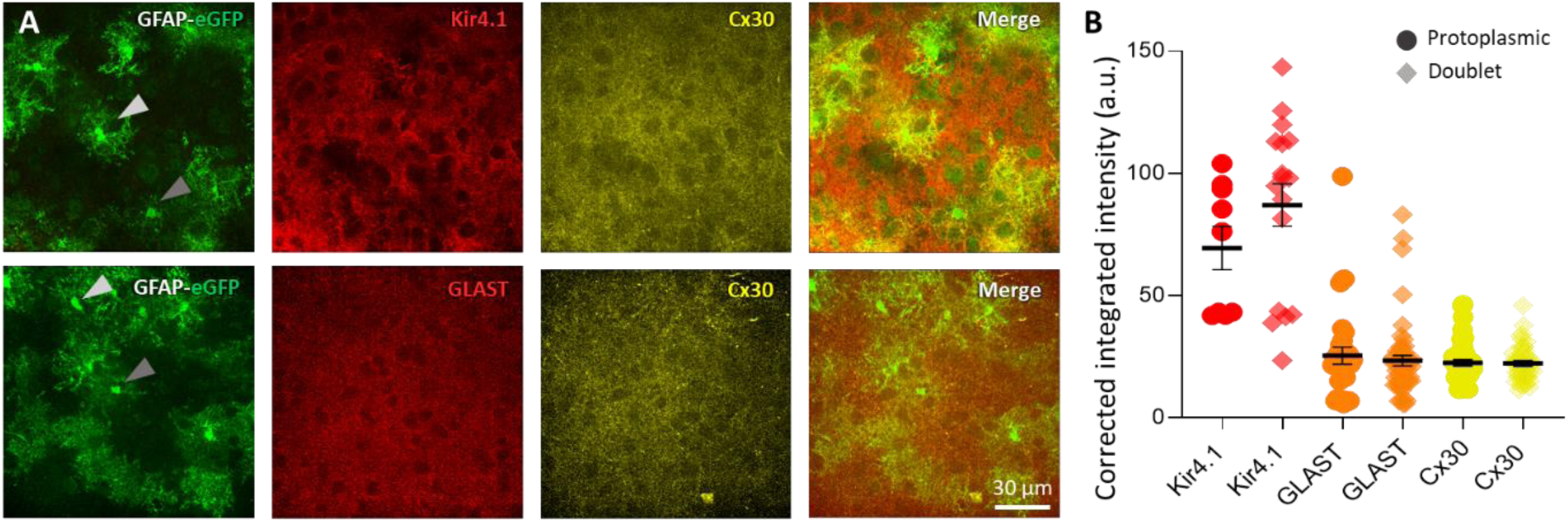
Molecular characterization of astrocytic subtypes: maturation status of doublet astrocytes and lineage identity of long-projection astrocytes. (A) Confocal images of GFP labeling (green), Kir4.1 (red, top) or GLAST (red, bottom), and Cx30 (yellow) labeling. The merges are shown in the right column. Gray arrowheads point to protoplasmic astrocytes (dark gray) or doublet astrocytes (light gray). (B) Quantification of corrected integrated intensity for each protein of interest in protoplasmic and doublet astrocytes, calculated as: Integrated Density within the ROI – (ROI Area × Mean Background Intensity). n=12 ROIs for GFP, Kir4.1, and Cx30, and n=10 ROIs for GLAST from 4 mice. Mann-Whitney, not significant.

### Long-projection Astrocytes Extend Their Processes Across Glial Territories and are restricted to the VLPO

To confirm the astrocytic nature of these long processes, we first examined the expression of the glial fibrillary acidic protein (GFAP), a canonical intermediate filament protein, frequently used as a marker of astrocytes. Immunostaining in wild-type C57Bl6 mice revealed strong GFAP labeling within these structures, which appeared relatively straight, unbranched, and up to twice the thickness of typical primary astrocytic processes (Figure S3A), supporting their astrocytic identity. To further assess their presence across different genetic backgrounds, we examined two transgenic reporter lines: Aldh1l1-eGFP, which labels the majority of astrocytes, and GFAP-eGFP, which marks a more restricted astrocyte population. At P30, long processes were consistently detected in both lines (Figure S3B-C).

These long processes are predominantly located near the pial border and oriented perpendicularly to it at this developmental stage.

**Figure S3.**
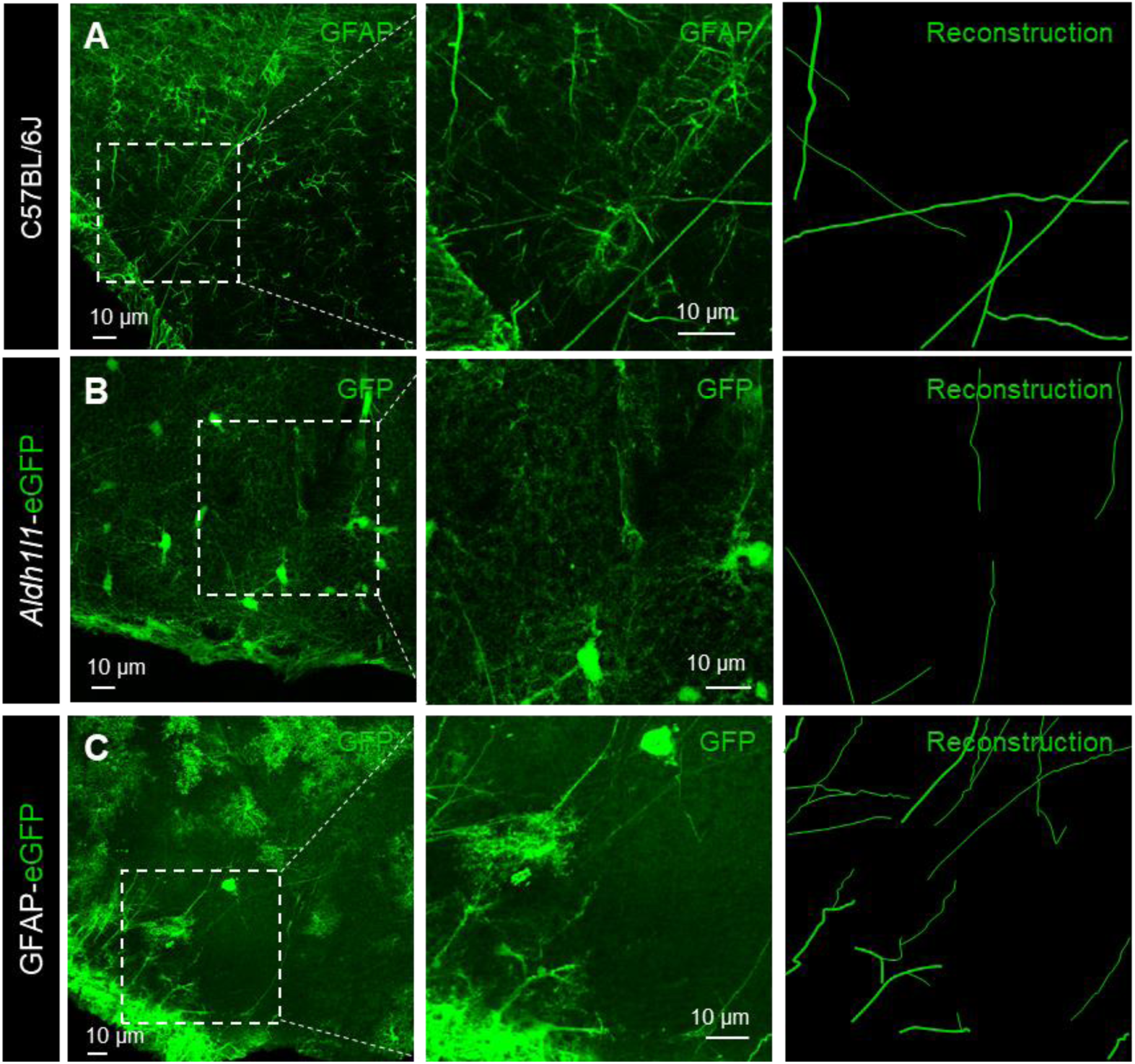
Long-projection astrocytes in the VLPO exhibit typical molecular profiles of astrocytes and are consistently observed across multiple mouse lines. (A) Confocal microscopy images from a C57BL/6J mouse, immunolabeled with GFAP (left). Magnified view of the dashed white square, highlighting the intricate long projection processes (middle). Reconstruction of these long projection processes (right). (B) and (C) Same as in (A), but in the Aldh1l1-eGFP (b) and GFAP-eGFP lines (C), respectively, with GFP immunolabeling in green (left and middle). The reconstructed images of these projections are presented in the right column for clearer structural representation. Note that long astrocytic processes are still observed, although they appear finer compared to the GFAP immunolabelling.

To determine whether these long astrocytic processes are specifically located within the VLPO, we performed GFAP immunolabeling in Gal-GFP transgenic mice. In this mouse model, sleep-promoting neurons of the VLPO that express galanin are endogenously labeled with green fluorescence, allowing precise anatomical delineation of the VLPO (Figure 5A–B). GFAP immunostaining revealed that long projections of astrocytes are restricted to the VLPO, supporting their regional specificity.

**Figure 5.**
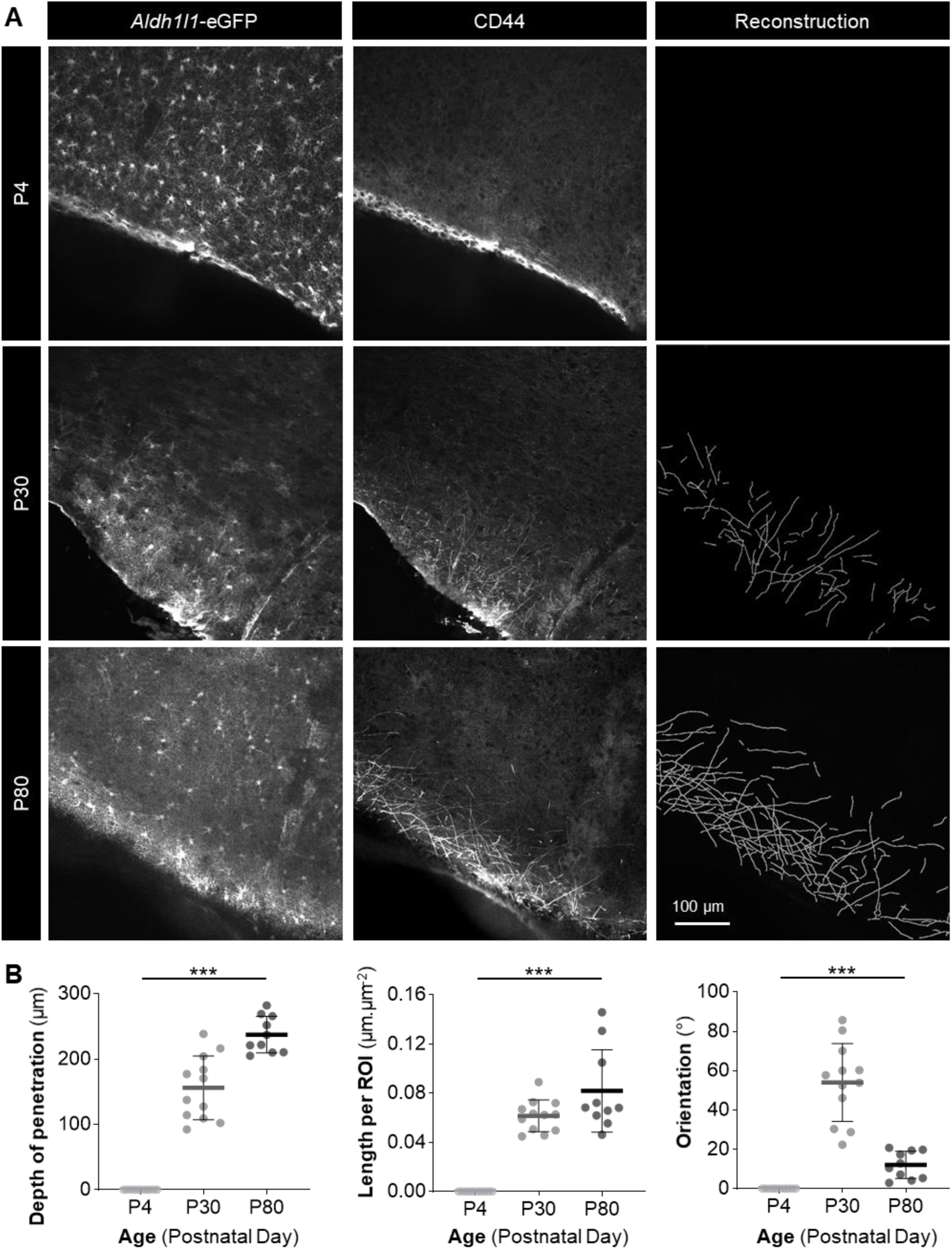
Long-projection astrocytic processes emerge during postnatal development and follow a depth-to-radial maturation trajectory. (A) Z-Stacks of 3 confocal microscopy images showing long-projection processes from an *Aldh1l1*-eGFP mouse at P4, P30, and P80, immunostained with GFP (left) and CD44 (middle). Reconstructions of their CD44^+^ processes are shown in the right column. (B) Measurements of the depth of penetration, length, and orientation relative to the pia of long-projection astrocytic processes through parenchyma (n = 10 at P4, n = 12 at P30, and n = 10 at P80). Kruskal-Wallis test. ***P < 0.001.

We next investigated how these long projections integrate into the spatial architecture of astrocytic domains and networks. Astrocytes are typically characterized by highly branched processes that tile the brain in non-overlapping domains. To investigate whether long-projection processes respect these spatial boundaries or, conversely, cross them without altering their trajectory, we performed GFAP immunostaining on GFAP-eGFP mice. This dual-labeling approach enables simultaneous visualization of long projections (shown in green) and the individualized astrocytic domains (shown in red for better visualization, Figure 5 C-E). Strikingly, our results revealed that long projection processes do not respect the conventional domain boundaries^48,49^, by extending their processes across neighboring territories, without altering their trajectory.

### Postnatal Emergence and Maturation of VLPO Long-Projection Processes

Numerous biological processes undergo significant changes during the first postnatal weeks, either developing further or regressing. For instance, delayed-formed astrocytes mature and develop more intricate structures, whereas the radial processes of radial glial cells diminish as the brain matures^50^. To determine whether the long-projection processes observed at P30 are transient, stable, or still maturing during postnatal development, we analyzed their characteristics at key postnatal stages: P4, P30, and P80. To selectively visualize these long processes and distinguish them from typical astrocytic arbors, we used the CD44 antibody. This marker was chosen based on the striking morphological similarity between VLPO long-projection astrocytes and the primate-specific interlaminar astrocytes described in the cortex, which are also known to express CD44^26,51^. Consistent with this hypothesis, we confirmed robust CD44 expression in VLPO long-projection processes. This molecular feature enabled us to reconstruct these structures in detail using Imaris software. The CD44 immunostaining was performed in Aldh1l1-eGFP mice, allowing us to visualize the entire astrocyte morphology while specifically highlighting the long projections (Figure 5).

At P4, long-projection CD44⁺ astrocytic processes are scarce (Figure 5). By P30, these processes begin to emerge, extending predominantly perpendicular to the pial surface and localizing primarily within the first 150 μm beneath the pial border. Then, by P80, their architecture becomes significantly more complex. They are notably denser, exhibit a multidirectional orientation, and span a much larger portion of the VLPO, suggesting continued maturation and spatial expansion throughout postnatal development (Figure 5).

To determine whether long-projection astrocytes preferentially terminate on specific elements, such as somata or blood vessels, we examined their termination at P150, when astrocytic maturation is complete, in the VLPO of Gal-GFP mice (Supplementary Figure S4). We traced 63 long processes to their final contact and found that 3.28 ± 2.28% terminated in close proximity to galaninergic neurons (Figure S4A), 14.75 ± 4.54% on a cell body (Figure S4B), 29.51 ± 5.84% near a blood vessel (Figure S4C), 31.15 ± 5.93% on another astrocyte (Figure S4D), and 21.31 ± 5.24% within the parenchyma without a clearly identifiable terminal structure (Figure S4E). Taken together, these findings indicate that VLPO long-projection astrocytes preferentially establish terminal contacts with other astrocytes and with blood vessels.

**Figure S4.**
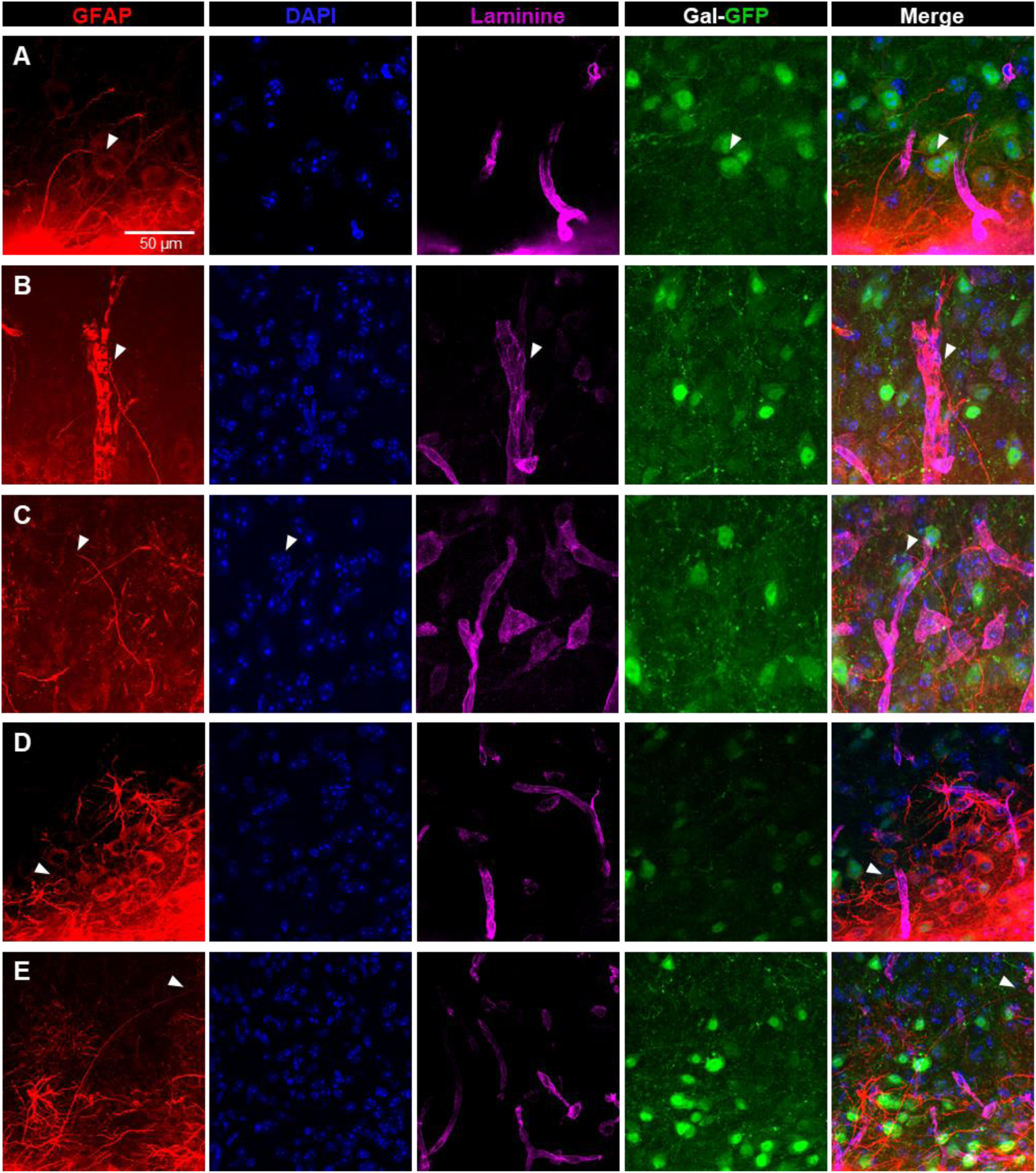
Long astrocytic processes preferentially terminate on other astrocytes and blood vessels. (A) Representative long process terminating in close proximity to a galaninergic neuron (GFP, green). (B) Representative long process terminating at a vessel (laminin, magenta). (C) Representative long process terminating on cell body (DAPI, blue). (D) Representative long process terminating near an astrocyte (GFAP, red) within the parenchyma without a clearly identifiable ending (GFAP, red). (E) Representative long process terminating in the parenchyma.

### Enhanced Ca^2+^ Signaling in VLPO Astrocytic Processes

Cellular morphology is a key determinant of functional properties^52^. We therefore investigated whether VLPO astrocytes exhibit functional features distinct from those of astrocytes in the cerebral cortex or hippocampus. As intracellular Ca^2+^ elevations constitute the principal readout of astrocytic activity, we characterized and compared spontaneous Ca^2+^ transients in astrocytes across these three regions, using genetically encoded calcium indicators (GECIs) in transgenic GFAP-CreERT2::GCaMP6f mice (Figure 6).

**Figure 6.**
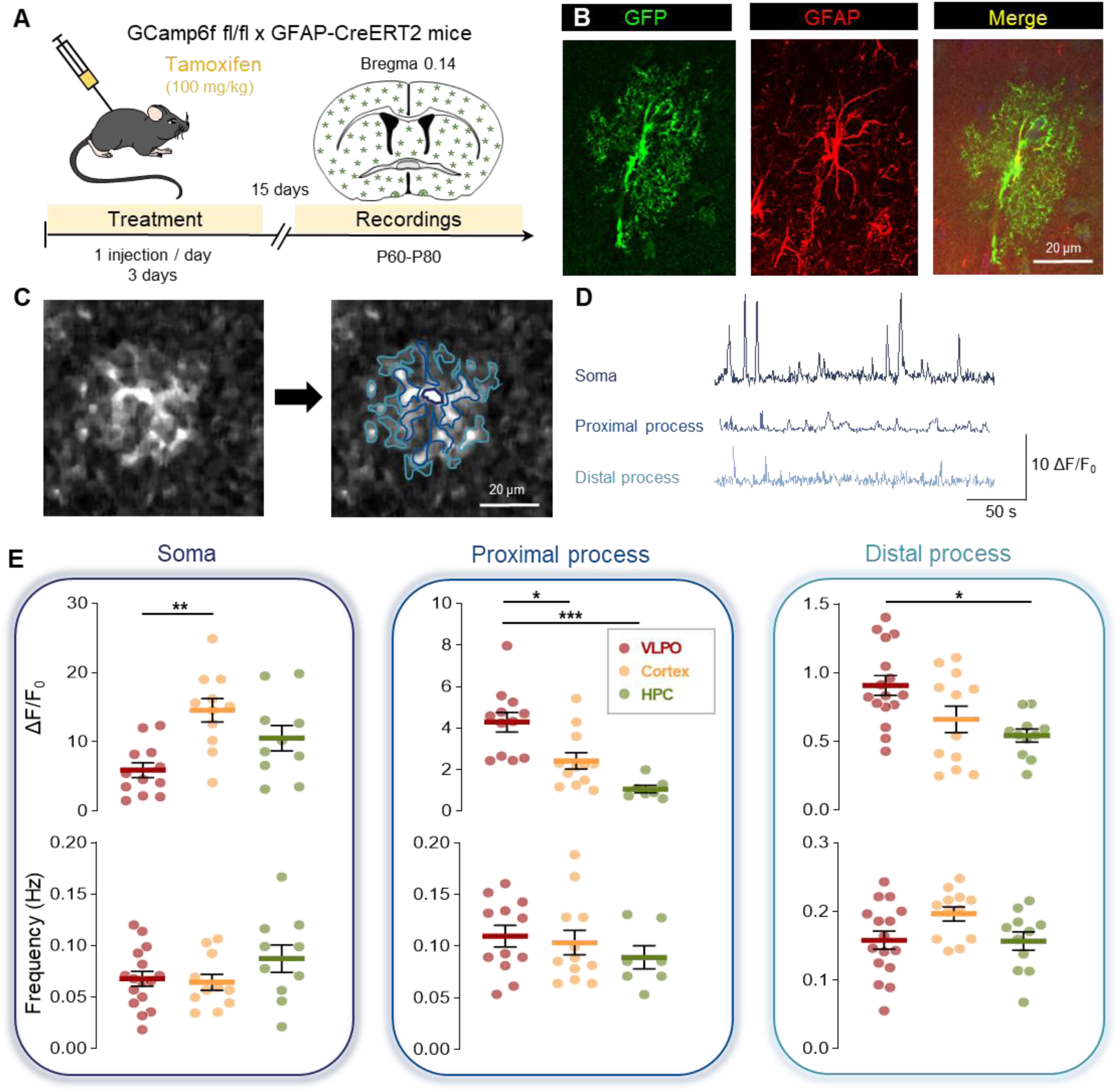
Spontaneous Ca^2+^ activity in VLPO astrocytic processes is enhanced compared to the cortical and hippocampal regions. (A) Design of the experimental protocol. (B) Confocal image of a GFAP immunostaining (red) in GFAP-CreERT2::GCaMP6f mice showing the specific expression of GCaMP6f (green) in astrocytes. (C) Two-photon image of calcic signals (left) and subdivision into three specific territories. (D) Typical traces of Ca^2+^ recordings in the three astrocytic territories. (E) Analyses of the signals. n = 9 astrocytes in the VLPO, n = 9 astrocytes in the cortex and n = 11 astrocytes in the hippocampus, from at least 3 different mice. One-way ANOVA, Tukey’s multiple comparisons test. ***P < 0.001, **P < 0.01, *P < 0.05.

Two-photon microscopy enables high-resolution imaging of astrocytic Ca^2+^ dynamics at both cellular and subcellular scales. However, this technique presents two main limitations: only astrocytes exhibiting spontaneous Ca^2+^ activity are detectable, and image acquisition is restricted to a two-dimensional plane. These constraints limit reliable morphological discrimination of long-projection astrocytes from protoplasmic astrocytes. Nevertheless, by imaging regions distant from the pial border, where long projection astrocytes are preferentially located, we were able to compare spontaneous Ca^2+^activity across protoplasmic and doublet astrocytes within the VLPO.

We first focused our analysis on protoplasmic astrocytes to enable a direct comparison with astrocytes from the cortex and hippocampus, thereby establishing a regional framework for subsequent intra-VLPO comparisons (Figure 6).

Across regions, we detected a clear compartment-specific reshaping of astrocytic Ca²⁺ signals. Somatic Ca²⁺ transients in VLPO astrocytes displayed a tendency for smaller amplitudes than those measured in cortical and hippocampal astrocytes. In contrast, Ca²⁺ events recorded in VLPO processes were markedly larger: both proximal and distal processes exhibited significantly higher event amplitudes compared with the corresponding compartments in cortex and hippocampus (Figure 6). Notably, these amplitude differences were not accompanied by changes in event frequencies, as mean Ca²⁺ event frequency remained comparable across the three regions for the soma, proximal processes, and distal processes.

Consistent with these regional features, we next examined whether astrocyte morphology within the VLPO is associated with distinct Ca²⁺ dynamics. Doublet astrocytes displayed clearly independent Ca²⁺ activity in each soma, indicating that the two closely apposed cells maintain functionally distinct signaling profiles (Movie S1). In addition, Ca²⁺ signals were observed to propagate along long-projection processes, confirming that these processes are functionally active compartments (Movie S2).

Quantitative analysis showed that doublet astrocytes display a higher Ca²⁺ event in the soma ∼+230%. In distal processes, event frequency tended to be increased in doublet astrocytes ∼35% (Figure S6). Together, these findings indicate that doublet astrocytes exhibit a compartment-specific upscaling of spontaneous Ca²⁺ signaling, consistent with a distinct functional organization compared with protoplasmic astrocytes.

**Figure S6.**
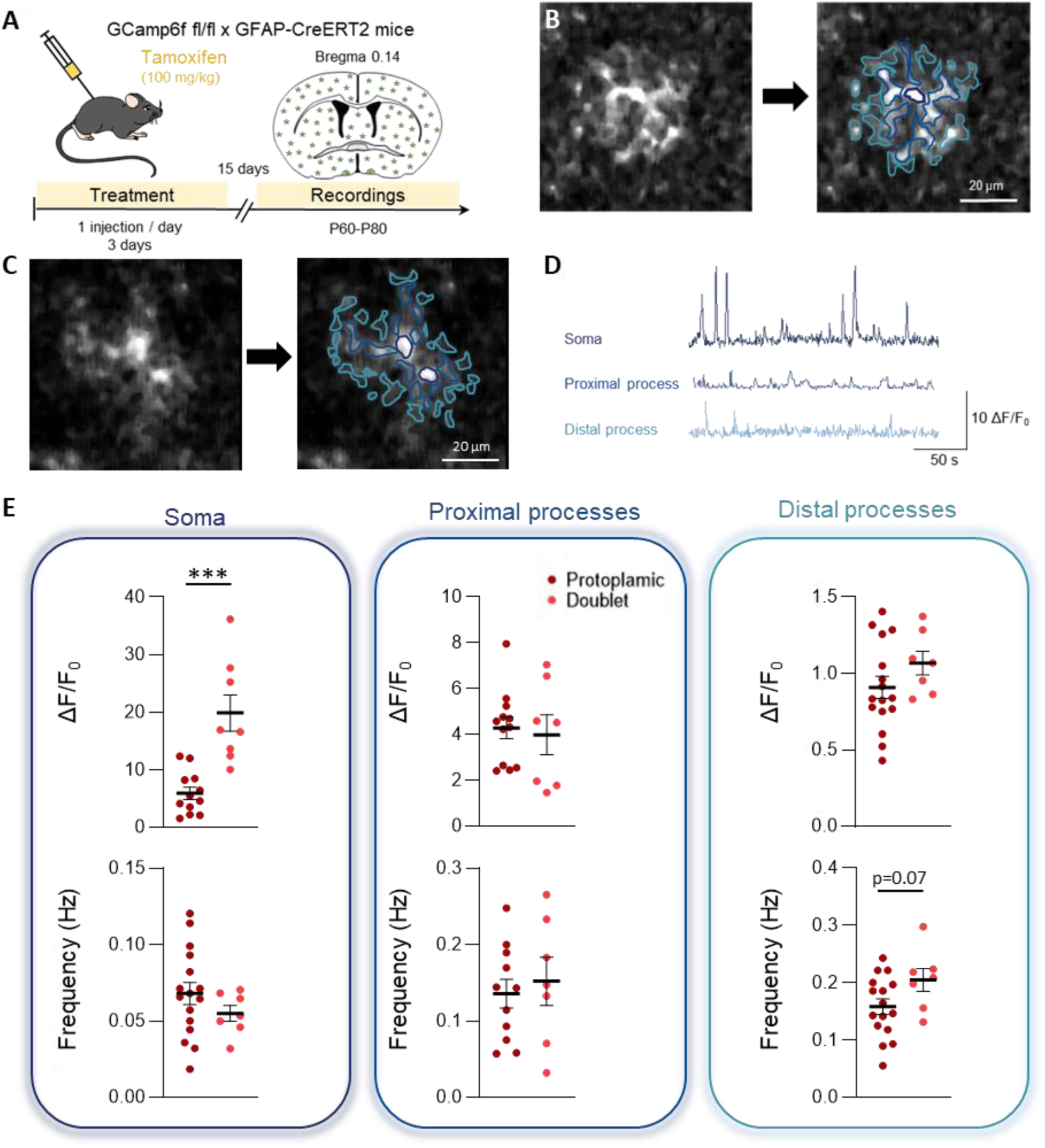
Astrocytic calcium signaling is enhanced in the soma, whereas event frequency tends to decrease in distal processes, in doublet astrocytes compared to protoplasmic astrocytes. (A) Design of the experimental protocol. (B) Two-photon image of calcic signals (left) in protoplasmic astrocytes and subdivision into three specific territories (right). (C) Two-photon image of calcic signals (left) in doublet astrocytes and subdivision into three specific territories (right). (D) Typical traces of Ca^2+^ recordings in the three astrocytic territories. (E) Analyses of the signals. n = 15 protoplasmic astrocytes in the VLPO, n = 8 doublet astrocytes in the VLPO, from at least 3 different mice. Unpaired t-test, Mann-Whitney test. ***P < 0.001.

Astrocytic Ca^2+^ signaling primarily relies on the inositol-1,4,5-trisphosphate receptor (IP3R) pathway. Local Ca^2+^ influx through the plasma membrane triggers further Ca^2+^ release from internal stores, mainly the endoplasmic reticulum and mitochondria, via IP3R activation. There are three isoforms of IP3Rs: IP3R1, IP3R2, and IP3R3, each exhibiting distinct activation and deactivation patterns. The expression and subcellular distribution of these isoforms vary across tissues and cell types^53^, contributing to their diverse functions and regulatory mechanisms. In astrocytes, the typical isoform transcript abundance is ITPR2 > ITPR1 >>> ITPR3, with ITPR3 mRNA potentially negligible^54^. Nevertheless, these transcripts are differentially regulated during development and across brain regions^54^ and they underlie distinct spatiotemporal Ca^2+^ dynamics^55^. To investigate whether subcellular differences in IP3R1 and IP3R2 distribution could account for the Ca^2+^ dynamics observed in the VLPO, we performed isoform-specific immunolabeling and quantitative analysis in astrocytic compartments of the VLPO, cortex, and hippocampus. Taking advantage of the GFAP-GFP mouse model, which enables subtype discrimination in the VLPO, we also compared IP3R expression among protoplasmic, doublet and long-projection astrocytes (Figure S5).

**Figure S5.**
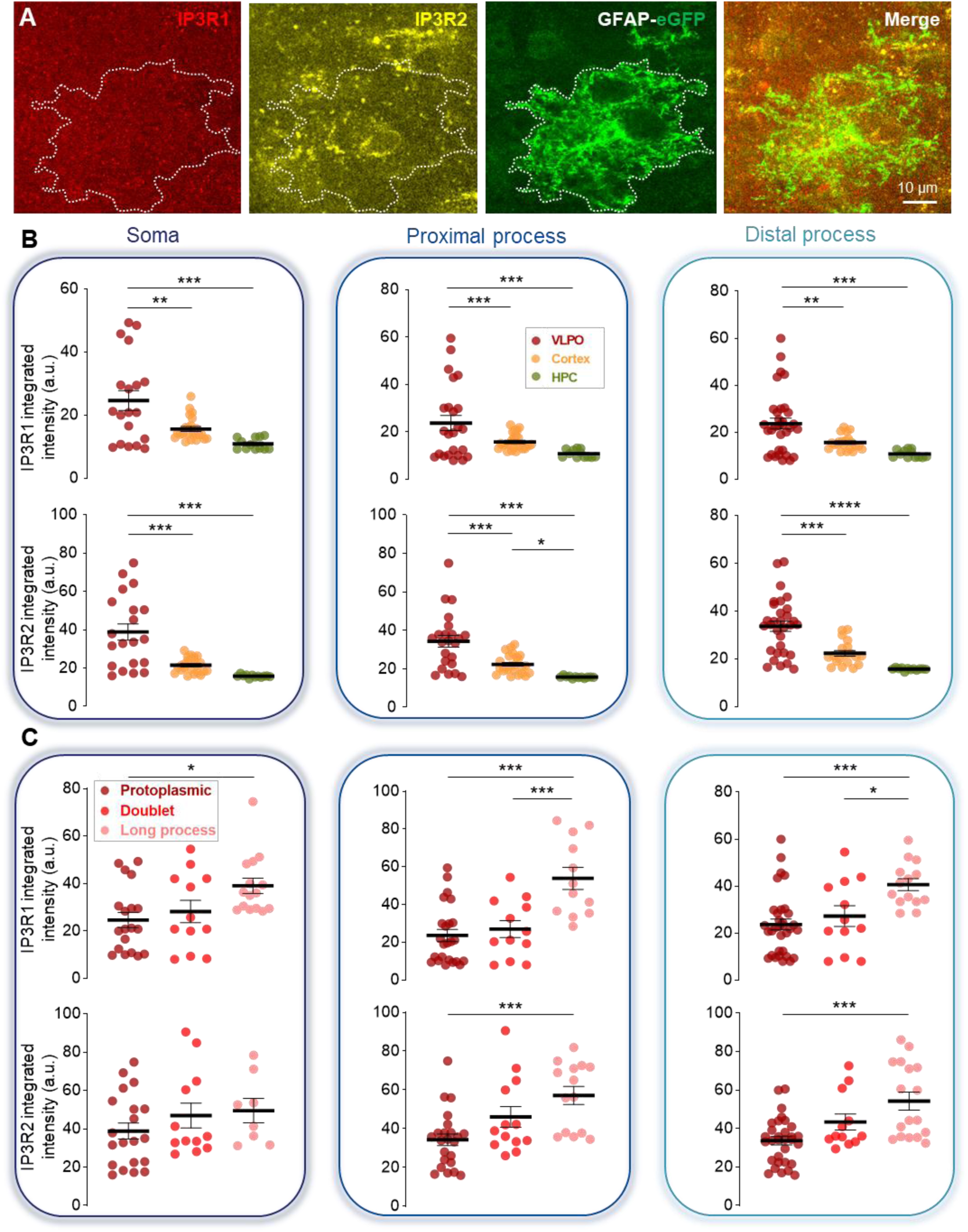
Differential Expression of IP3R1 and IP3R2 Across Brain Regions and Astrocyte Subtypes. (A) Representative confocal image of the immunostaining for IP3R1 (red) and IP3R2 (yellow) from GFAP-eGFP mice (green). (B) Quantification of the IP3R1 (top) and IP3R2 (bottom) integrated intensity within the soma (left), proximal process (middle) and distal microdomain (right), across the VLPO (red), cortex (yellow), and hippocampus (green). (C) Same as in B, specifically comparing the 3 astrocyte subtypes of the VLPO. n=18 ROIs in 5 mice for the VLPO; n=22 ROIs in 4 mice in the cortex and n=16 ROIs in the HCP. One-way ANOVA, Tukey’s multiple comparisons test. ***P < 0.001, **P < 0.01, *P < 0.05.

We observed that both IP3R1 and IP3R2 isoforms are significantly enriched in the VLPO across all astrocytic subcellular compartments compared to the cortex and hippocampus (Figure S5). Notably, their expression is further elevated in long-projection astrocytes. This molecular enrichment likely underlies the heightened Ca^2+^ signaling observed in these cells, as elevated levels of IP3 receptors enhance the cell’s capacity for intracellular Ca²⁺ release from internal stores. Such a mechanism may support more robust, widespread, or sustained Ca^2+^ dynamics, contributing to the unique functional properties of VLPO astrocytes.

### VLPO Astrocytes form a Highly Connected Functional Network

Astrocytes are organized into networks whose sizes are tightly optimized to regulate neuronal networks appropriately^56^. However, these astrocytic networks’ functional activity and organization remain poorly studied, although they can reveal interesting differences in network dynamics between distinct brain regions. To gain insight into the functional organization of astrocytic networks, we analyzed spontaneous Ca^2+^ activity from the VLPO, cortex, and hippocampus using the AstroNet MATLAB toolbox^42^. This approach enables the identification and tracking of activated pathways between successively activated astrocytes, the reconstruction of connected astrocyte networks, and the creation of functional connectivity graphs (Figure 7A). Subsequently, we analyzed these graphs to extract various network parameters (Figure 7B-I).

**Figure 7.**
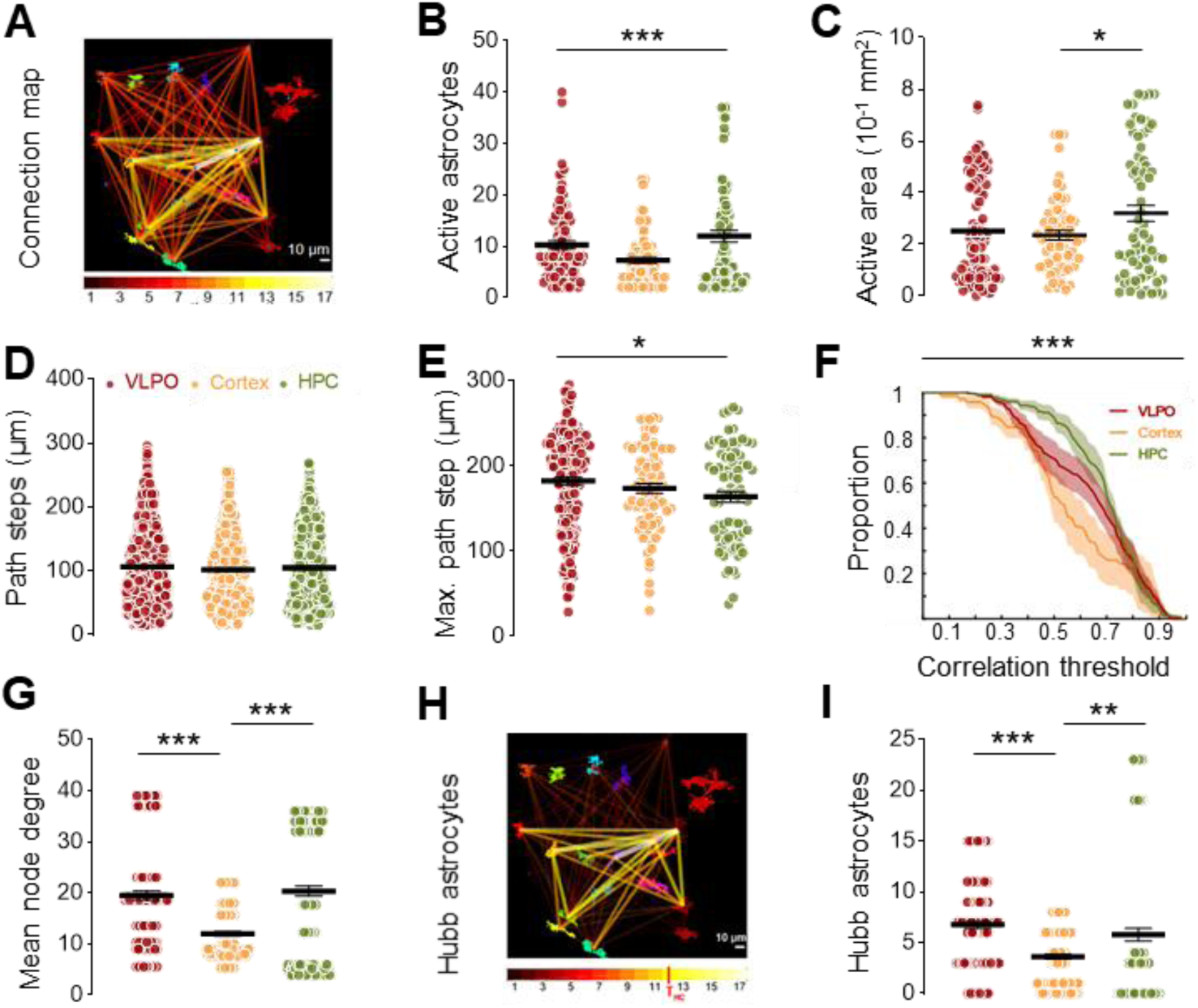
Functional connectivity of astrocytic networks in the VLPO, cortex, and hippocampus. (A) Analysis of the astrocyte functional network allows the reconstruction of local connectivity patterns, as shown in this reconstructed graph, which is color-coded according to different connectivity levels: strong (yellow) to light (dark red). (B) Number of active astrocytes in a recording. (C) Surface of the convex hull of the active area per recording. (D) The path steps correspond to the distances between each pair of consecutively activated astrocytes. (E) Maximal path step per recording. (F) Pairwise correlation matrix between all astrocyte time-series enables correlation analysis across the three regions. A faster decay reveals a less correlated network. (G) Mean node degree in reconstructed graph. (H) Functional network of hub astrocytes. (I) Number of hub astrocytes. Two-way and one-way ANOVA, Kruskal-Wallis, and Dunn’s and Tukey’s multiple comparisons test. ***P < 0.001, **P < 0.01, *P < 0.05.

Functional analysis of astrocytic networks revealed that a higher proportion of active astrocytes is present in the hippocampus compared to the cortex, with the VLPO displaying intermediate levels (Figure 7B). A similar trend was observed for the active area, reflecting broader spatial activation in the hippocampus (Figure 7C). While the average length, defined as the distance between consecutively activated astrocytes, did not significantly differ across the three regions (Figure 7D), a more detailed spatial analysis revealed that the VLPO contains significantly longer maximal steps (Figure 7E). These maximal steps correspond to the longest individual distances over which calcium signals propagate between two consecutively activated astrocytes during the 30 s recording. Such extended connections are likely enabled by the presence of long-projection astrocytes in the VLPO.

To get further insights into the functional organization of these networks, we estimated the overall connectivity by the pairwise Pearson correlation between active astrocytes. We generated a mean correlation curve for each region by computing the proportion of astrocytes having at least one correlation with another astrocyte stronger than a correlation threshold increasing from 0 to 1 (Figure 7F). The closer the curve is to the top left corner, the more globally connected the astrocyte network. Our results revealed that the VLPO (red) exhibits a global intermediate level of connectivity, greater than that observed in the cortex (yellow), but lower than in the highly connected hippocampus (green). Interestingly, the bump in the VLPO curve indicates the presence of more highly connected astrocytes than in the cortex.

Hence, we counted the number of direct neighboring astrocytes for each astrocyte in a given network: the node degree, and found that the mean node degree in the VLPO is comparable to that of the hippocampus and almost double what is observed in the cortex (Figure 7G). Furthermore, we can see that the VLPO (red) exhibits a small cluster of astrocytes with a node degree of around 38-40, which is even higher than those in the hippocampus, which range up to ∼36 (green).

To further evaluate astrocytic connectivity, we counted the number of astrocytes exhibiting high connectivity. The nodes of these highly connected subgraphs can be considered hub astrocytes, i.e., astrocytes that participate in almost all activation events within the local network. We choose to count astrocytes that participate in at least 60% of events (Threshold THC = 0.6 ∗ 2N for undirected graphs, where 2N is the maximal strength of an 8-edge). We found that the VLPO has more hub astrocytes, indicating more internal connections.

## Discussion

Despite the crucial role of astrocytes in regulating neuronal activities, VLPO astrocytes have only very recently begun to be investigated. This work is the first to report the identification of three distinct astrocytic subtypes in the VLPO: protoplasmic, doublet, and long-projection astrocytes. These findings highlight the morphological and functional heterogeneity of astrocytes in this region and suggest their potential roles in sleep regulation.

### Morphological Specialization and Regional Diversity

Protoplasmic astrocytes are the most abundant glial cell type in the VLPO, consistent with their prevalence in gray matter across both rodent and primate brains^44^. However, these cells exhibit region-specific differences^57^. In the VLPO, protoplasmic astrocytes display smaller domains than those in the cortex and hippocampus, and arborization complexity is lower. These results are consistent with previous findings, which show that astrocytic volumes vary among brain regions and across species. In rodents, these volumes range from ∼15 000 to 80 000 mm ^3^ ^18,23,48,58–60^. Such differences underscore the region-specific specialization of astrocytes, potentially tailored to the VLPO’s role in sleep regulation.

Astrocytic doublets in the VLPO are defined by their spatially constrained domains and tightly apposed somata. Our findings indicate that the elevated and sustained astrocytic proliferation observed in the VLPO is a key driver of doublet formation. This contrasts with the sharp decrease in astrocytic proliferation during cortical development, notably after the first postnatal week, and which is almost entirely halted by P21^61,62^. Although the VLPO also exhibits a decline in proliferation by P30 and P80, its proliferation rate remains substantially higher than that of the cortex, resulting in ∼20 times more doublets in the VLPO than in the cortex by P80. Postnatal astrocyte generation primarily involves the local proliferation of “pioneer” astrocytes, which divide symmetrically^62^. The increased spatial constraints within the VLPO progressively limit the separation of newly divided astrocytes^63^, ultimately leading to a higher prevalence of doublets.

Long-projection astrocytes, previously presumed to be exclusive to human and primate brains^27,63^, have now been identified in the mouse VLPO. Although rare protruding processes have been observed in the rat hippocampus, these processes also exhibit no branching, but are typically much shorter, only a few micrometers in length, and generally retract or degenerate by one month of age^49^. Bushong’s observations also revealed that these processes, as in the VLPO, are strongly labeled by GFAP immunostaining, indicating their richness in cytoskeletal proteins. However, unlike the persistent development of long-projection astrocytes in the VLPO, these hippocampal processes are typically retracted or degenerate by one month of age.

In humans, long-projection astrocytes include two types: interlaminar astrocytes and varicose-projection astrocytes. Interlaminar astrocytes have either a soma located in the first cortical layer and processes extending toward the deep layers or a soma located in layers 5-6^27,64^. In contrast, varicose-projection astrocytes, which exhibit varicosities along their long processes, are rare, reside in the deep layers of the cortex, and are found exclusively in hominoid brains^64,65^. VLPO long-projection astrocytes lack varicosities, suggesting they more closely resemble interlaminar astrocytes.

In monkeys, interlaminar astrocytic fibers are initially short and sparse during early postnatal development, but gradually increase in length and number with maturation^66^. This trajectory parallels the development of VLPO long-projection astrocytes. Furthermore, similar to cortical long-projection astrocytes, those in the VLPO interact with blood vessels, perivascular astrocytes, neurons, and protoplasmic astrocytes. This connectivity suggests a significant role in integrating vertical and horizontal information flow within the VLPO^26^.

### Functional Implications of Astrocytic Heterogeneity

Morphological heterogeneity among protoplasmic astrocytes is aligned with their distinct calcium signaling profiles across brain regions. In the VLPO, astrocyte calcium events in proximal and distal processes are notably more pronounced than in cortical and hippocampal astrocytes. This heightened activity is likely supported by the elevated density of IP3R1 and IP3R2. Accordingly, a high density of IP3R2 is associated with elevated Ca^2+^ release kinetics^67^. This amplified signaling capacity likely reflects an adaptation to the VLPO’s pivotal role in regulating sleep-wake cycles, where astrocytic Ca^2+^ activity has been implicated in sleep regulation through the release of sleep-promoting molecules, such as adenosine, and the modulation of neuronal activity^68^. Astrocytic networks indeed contribute to the coordination and regulation of neuronal network dynamics, influencing local and global brain activities^69^. These findings underscore the intricate interplay between astrocytic Ca^2+^ signaling and the functional demands of the VLPO in sleep regulation.

Delayed astrocytic proliferation in the VLPO, responsible for the numerous formations of doublets, may be critical for synapse formation, maturation, and pruning^70,71^, as well as for metabolic support^71^ during a developmental window. This prolonged proliferation could support the significant maturation of sleep patterns that occurs during development.

From an evolutionary perspective, sleep likely enhances cognitive capacities in species^72^, while simultaneously exposing them to increased vulnerability to predators due to reduced vigilance. Evolutionary pressure may therefore have favored an increased amplitude of Ca^2+^ responses in the VLPO, optimizing the efficiency of underlying neural networks^73,74^ and facilitating rapid transitions between vigilance states^75^. Functional connectivity analysis further revealed that VLPO astrocytes form highly integrated functional networks. The presence of hub astrocytes suggests a specialized architecture that supports robust communication within the VLPO. The presence of long-projection astrocytes may enhance the efficiency of neuronal network communication in the VLPO by facilitating rapid and widespread propagation of astrocytic signals, even beyond the analyzed areas in this study. Their presence in the VLPO, a region integral to sleep regulation, suggests that evolutionary pressures may have driven the emergence of specialized astrocytes to meet the demands of complex brain functions.

Intralaminar astrocytes in hominids are hypothesized to participate in immune defense, regulate synapse formation and function, and modulate the glymphatic system and water homeostasis at the blood-brain barrier and the pia interface. Their unique microRNA expression profile suggests a specialized “first-line defense” role against external insults^76^. Moreover, their proximity to synapses and the expression of aquaporin-4 (AQP4) indicate a potential involvement in maintaining synaptic homeostasis and fluid regulation^64^. Further studies are required to explore whether VLPO long-projection astrocytes share similar properties.

## Conclusion

The identification of three distinct astrocytic subtypes in the VLPO provides new insight into astrocyte heterogeneity. More than a structural observation, this diversity suggests specialized roles in the modulation of neuronal circuits and sleep maturation. Future research should aim to elucidate the molecular mechanisms driving this specialization and explore their relevance in the physiological regulation and pathological disruption of sleep.

## Supporting information

Supplemental Moovie 1

Supplemental Moovie 2

## Acknowledgments

We thank all the members of the animal house facility from the Collège de France and of the laboratory “Neuroglial Interactions in Cerebral Physiology and Pathologies” for the discussions on this work. The authors particularly thank Isabelle Arnoux for genotyping the GCamp6f mice.

## Author Contributions

Conceptualization: A.R. and Methodology, L.Z., D.H, Q.P., A.R.; Formal Analysis, F.C.B., A.R., and L.Z.; Investigation, F.C.B.; Writing – Review & Editing, A.R.; Critically revised the manuscript, all authors; Visualization, F.C.B., A.R. and L.Z.; Funding Acquisition, N.R.; Resources, K.L..; Supervision and project administration, A.R.

## Declaration of generative AI and AI-assisted technologies in the writing process

During the preparation of this work, the authors used Chat-GPT to improve grammar and English writing. After using this tool/service, the authors did not accept all the suggested changes and reviewed the content as needed and take full responsibility for the content of the published article.

## Supplemental information

Figures S1–S4

Video S1-S2

