## Supplementary figures and images for "Long-projection astrocytes challenge canonical territorial organization in the sleep-promoting VLPO"

### Supplemental Moovie 1

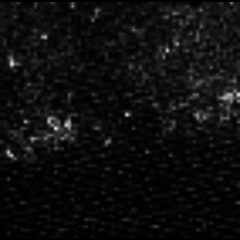

### Supplemental Moovie 2

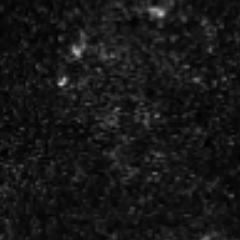
